# Historical biogeography of New World Killifishes (Cyprinodontiformes: Funduloidea) recapitulates geographical history in the Gulf of México watershed

**DOI:** 10.1101/2023.09.07.556706

**Authors:** Sonia G. Hernández, Christopher W. Hoagstrom, Wilfredo A. Matamoros

**Affiliations:** Programa de Licenciatura, Instituto de Ciencias Biológicas, Universidad de Ciencias y Artes de Chiapas. Libramiento Norte Poniente No.1150, Colonia Lajas Maciel, C.P. 29039. Tuxtla Gutiérrez, Chiapas, México; Maestría en Ciencias en Biodiversidad y Conservación de Ecosistemas Tropicales, Instituto de Ciencias Biológicas, Universidad de Ciencias y Artes de Chiapas, Tuxtla Gutiérrez, Chiapas, México; Department of Zoology, Weber State University, 1415 Edvalson, Ogden, Utah, USA

**Keywords:** barrier displacement, freshwater–fish paradox, Cyprinodontidae, Cubanichthyidae, Fundulidae, Goodeidae, Profundulidae, reciprocal illumination, speciation pump, speciation rate

## Abstract

We reconstructed the phylogenetic relationships of superfamily Funduloidea with a synthesis of its biogeographic history. We used DNA sequences from five genes for 135 species, with four fossil calibrations, to generate a time-calibrated phylogeny. We estimated diversification rates, ancestral areas (Nearctic or Neotropical), and ancestral habitat (coastal or upland), for each node. Our results suggest that Funduloidea originated in the Late Cretaceous and diversified from Late Paleocene to present at a uniform rate, except Cyprinodontidae expressed an accelerated rate of speciation ~11.02 Ma. Neither viviparity, marine-to-freshwater transition, consistently accelerated speciation. Funduloidea has a coastal origin, but invaded inland many times. Funduloidea phylogeny indicates, sea-level falls isolate coastal populations, but increase island accessibility and climatic cooling facilitates invasions of temperate species into the tropics. For continental lineages, ancient river drainages accord with lineage distributions, including enigmatic disjunctions in Goodeidae and *Fundulus*. Niche shifts occurred from estuaries to open coasts and from forests to grasslands. Antiquity, adaptability, and dynamic geography can explain Funduloidea diversity. Combined environmental and phylogenetic data unveil the history of the Gulf of México watershed. Phylogeny suggests there was diversification by barrier displacement and coastal speciation pump. Overall, speciation time, transitions to freshwater, dispersal, vicariance, adaptive radiation, and viviparity contributed to total diversification.

## INTRODUCTION

Historical biogeography attempts to explain how past events produced current species distributions (Gutiérrez-García & Vázquez-Domínguez, 2013; Mastretta-Yanes et al., 2015). Within freshwater fishes, evolution of river-drainage configurations can mediate lineage diversification (e.g., Sidharthan et al., 2020; Cassemiro et al., 2023). Thus, studying phylogenetics of freshwater fishes goes hand in hand with studying geographic evolution. This relation of geology to phylogeny has been termed reciprocal illumination for the ability of these two fields to work together to solidify understanding of species diversification and Earth history (Santos & Capellari, 2009; Waters et al., 2019).

Also of interest in historical biogeography are key events that facilitate diversification. For example, although freshwater fishes appear to have high biodiversity compared to marine fishes (i.e., freshwater-fish paradox, Bloom et al., 2013; Tedesco et al., 2017; McDermott, 2021), recent evidence indicates this is not a general phenomenon and that only certain clades of freshwater fishes (Rabosky, 2020) or certain freshwater habitats (Miller, 2021) sustain accelerated speciation rates. Factors predicted to facilitate lineage diversification include ecological adaptability (Whitehead, 2010; Foster & Piller, 2018), viviparity (Helmstetter et al., 2016), adaptive radiation (Martin & Wainwright, 2011), and time-for-speciation (García-Andrade et al., 2021). Further, the process of barrier displacement can generate species diversity through fragmentation and recombination of fish populations (Albert et al., 2018). This can take various forms, such as river-drainage rearrangements mediated by tectonics, volcanism, and climate (e.g., Domínguez-Domínguez et al., 2010; Hoagstrom & Osborne, 2021), river captures in quiescent landscapes mediated by fluvial geomorphology (Albert et al., 2018; Val et al., 2022), and alternating sea-levels (Dolby et al., 2018). Cyclical processes like sea-level fluctuation may have reciprocating effects, producing a speciation pump (April et al., 2013; Owens, 2015). A phylogenetic synthesis paired with a review of geological and environmental evidence can identify instances in which these various processes have likely occurred.

Here, we investigate the biogeography of superfamily Funduloidea (Cyprinodontiformes, *sensu* Costa, 1998), a northern clade of New World killifishes that historically included Fundulidae, Goodeidae, and Profundulidae. However, recent phylogenomic studies indicate Cyprinodontidae is sister to Fundulidae (i.e., nested within Funduloidea), while the enigmatic Cubanichthyidae also belongs somewhere within the group (Ghezelayagh et al., 2022; Piller et al., 2022). Funduloidea is interesting because many processes associated with diversification are present. For instance, presence of tropical and temperate lineages implies there were niche shifts between realms. Similarly, presence of coastal and continental lineages implies there were shifts between marine and freshwater ecosystems (Whitehead, 2010). Funduloids also display adaptive radiations (Richards et al., 2021), viviparity (Helmstetter et al., 2016), and coastal dispersal (Dolby et al., 2018). A phylogenetic analysis of this superfamily provides a chance to put these varied processes into a broadened evolutionary context to better understand diversification in fishes.

In this study, we synthesize the biogeography of Funduloidea with an emphasis on early branching, using a newly generated phylogenetic tree. To begin, we look for instances of change in diversification rate to identify evolutionary steps where speciation rate changed (Helmstetter et al., 2016). Next, we conduct ancestral-areas analysis at a coarse scale (Neotropical versus Nearctic) to identify instances of biome switching. Third, we locate transitions from coastal to upland habitats. Fourth, we synthesize these findings with a review linking major branching events within Funduloidea to associated geological and environmental histories. We explore evidence for three factors that are expected to increase speciation rate, (1) adaptive radiation (Martin & Wainwright, 2011), (2) viviparity (Helmstetter et al., 2016), and (3) transition to freshwater (Bloom et al., 2013), and we assess whether speciation is faster in freshwater. This synthesis advances evolutionary studies within the superfamily, provides insight into broader evolutionary hypotheses, and illuminates how environmental and geological factors like sea-level fluctuation, river-drainage reorganization, and niche shifts may influence diversification and inland invasion by a euryhaline, coastal taxon.

## MATERIALS AND METHODS

### Sequence data

We constructed a Funduloidea phylogeny with data from GenBank (Supporting information, Table S1), including 135 species from a total of 160 recognized species. Outgroups were Atheriniformes (*Atherinomorus stipes, Atherinella alvarezi*) and Beloniformes (*Petalichthys capensis, Hemiramphus balao*). Data included two nuclear genes (recombination activating gene (RAG1), glycosyltransferase gene (Glyt)) and three mitochondrial genes (cytochrome b gene (Cytb), cytochrome oxidase 1 gene (COI), and NADH dehydrogenase subunit 2 gene (ND2)) (Table S1). A molecular data matrix was generated for each gene and individually examined and visualized in MEGAX v10.0.5 (Kumar *et al*., 2018). Sequences were aligned separately under default settings in the MUSCLE (Edgar, 2004) option in MEGAX. Each alignment was visually inspected, trimmed to equivalent sizes, and concatenated into a single matrix (5189 bp, 139 terminals; Supporting information, File S1) in Mesquite v3.61, Maddison & Maddison, 2016). For each gene, we performed a search for adequate nucleotide substitution models in Jmodeltest v2.1.10 (Darriba *et al*., 2012).

### Phylogenetic analyses

We performed a phylogenetic analysis under Bayesian inference in the program Mr. Bayes v3.2.7 (Ronquist *et al*., 2012) using the Markov chain Monte Carlo (MCMC) algorithm, run at ten million generations, sampling every 1000 iterations over four simultaneous runs. The analysis was verified to reach 0.01 average standard deviations of split frequencies. Subsequently, convergence was corroborated through Tracer v1.7.1 (Bouckaert *et al*., 2019). The 50% majority rule tree was visualized in FigTree v1.4.4 (Rambaut, 2010). The analysis was performed in the CIPRES v3.3 portal (Miller *et al*., 2010).

### Divergence-time estimations

We calculated divergence-time estimates to establish a temporal framework (Rutschmann, 2006). We used BEAST2 v2.6.3 (Bouckaert *et al*., 2019) to produce a time-calibrated phylogeny under Bayesian inference. The results of jModeltest v2.1.2 suggested the GTR model of molecular evolution was most appropriate for each locus (Cytb, COI, ND2, Glyt, RAG1). We opted for a relaxed-clock model assuming different substitution rates under the Speciation-Birth-Death model, which assumes constant speciation and extinction rates (Drummond *et al*., 2006). We calibrated the tree with four fossils used in previous studies, with priors fitted to a lognormal distribution (Table 1). Ghedotti & Davis (2017) vetted three of these fossils from *Fundulus*. A fourth fossil, †*Empetrichthys erdisi*, comes from the Posey Canyon shale, Peace Valley formation of the Ridge Basin, southern California (Uyeno & Miller, 1962). Uyeno & Miller (1962) cited this as a Pliocene deposit (see also Smith *et al*., 2002; Webb, 2020). However, the Peace Valley Formation is actually Late Miocene, 8.4-5.0 Ma (Crowell, 2003), which we follow here (Table 1). The Posey Canyon shale member, which holds †*E. erdisi*, is in the upper-middle Peace Valley formation (Link, 2003). We excluded two fossils used in recent studies. First, although Helmstetter *et al*. (2016) included “†*Cyprinodon breviradius*”, we did not because its identity and age are unclear (Smith *et al*., 2002; Echelle & Echelle, 2020). Second, although Webb *et al*. (2004) and Pérez-Rodríguez *et al*. (2015) included †*Tapatia occidentalis*, their phylogenetic placements were inconsistent. Upon review, we chose to exclude it because its position within Goodeidae is unknown (Smith, 1980; Guzmán, 2010). The divergence-time estimation analysis ran the Markov chain Monte Carlo (MCMC) for 50 million generations, sampled every 1000 generations in BEAST2 v2.5 (Bouckaert *et al*., 2019). To evaluate convergence and effective sample size (>200), we employed Tracer v1.7.1 (Rambaut *et al*., 2018) and used LogCombiner v2.6.3 (Bouckaert *et al*., 2019) to combine the replicates, discarding the first 25% of burns for each execution. Finally, we used TreeAnotator v2.6.3 (Bouckaert *et al*., 2019) to obtain a maximum-clade-credibility tree.

**Table 1.**
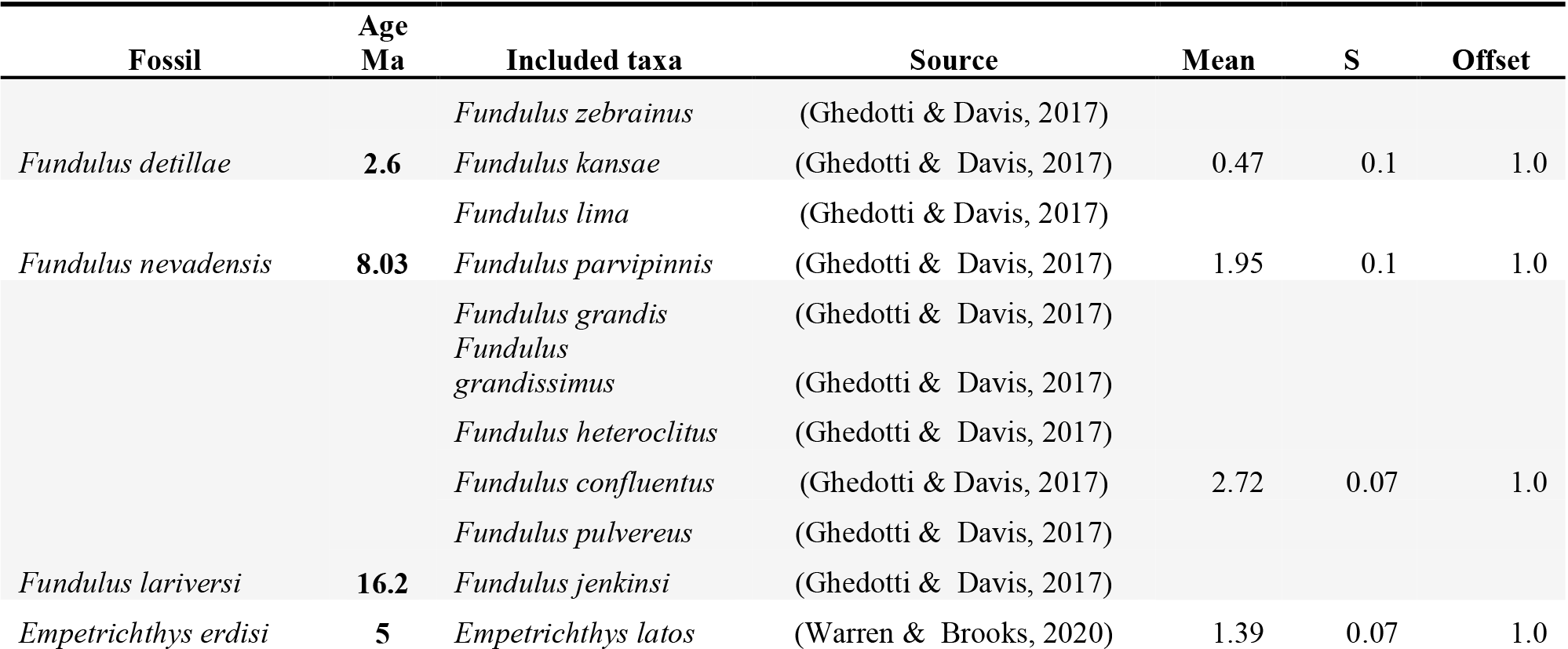
Fossils used in molecular dating analysis as calibration points in chronological order. Included taxa at each calibration point and prior parameters: mean, standard deviation (S) and offset. Fossils are from Ghedotti & Davis (2017) and Uyeno & Miller (1962). Dates for *Empetrichthys erdisi* are for the Peace Valley formation (Ridge Basin, Southern California; Ensley & Verosub, 1982), which includes the Posey Canyon shale (Link, 2003), where E. erdisi occur (Uyeno & Miller, 1962). From each age range, our analysis used the youngest age (in bold) as a hard minimum (Parham et al., 2012).

### Diversification-through-time

To estimate diversification rates, we used the BAMM v2.5.0 R package (Rabosky *et al*., 2014), which estimated speciation and extinction rates from the BEAST2 tree. To acquire parameters for analysis, we generated prior values in the R package BAMMtools 2.0 (Rabosky *et al*., 2014), using setBAMMpriors. The file obtained was then executed in BAMM. We ran four independent chains with five million generations, sampling every 1,000 to ensure convergence. Subsequently, we corroborated convergence of the log-likelihood effective sample sizes and stationarity using BAMMtools with the R package coda v016-1 (Plummer *et al*., 2006). We discarded the first 10% of the samples as burnin and calculated diversification-rate-shift settings with 95% credibility using the CredibleShifSet function. To obtain the best change configuration with maximum posterior probability, we used the getBestConfiguration function. We plotted speciation rates over time with the plotRateThroughTime function to generate lineage-through-time (LTT) plots using ‘PlotRateThroughTime’ for each family (excluding Cubanichthyidae, which has only two species).

### Ancestral-areas reconstruction

Reconstruction of ancestral areas infers likely distributions of ancestors based on contemporary distributions and phylogenetic relationships (Joy *et al*., 2016). We reconstructed ancestral ranges in RASP v4 (Yu *et al*., 2020) using six statistical models, including (1) Statistical Dispersal-Vicariance (DIVALIKE) in which speciation by vicariance is given primacy while dispersal and extinction are minimized (Yu *et al*., 2010); (2) Dispersal Extinction and Cladogenesis (DEC) which allows more ways for widespread ranges to be subdivided and inherited than just by vicariance (Beaulieu *et al*., 2013); and (3) BAYAREALIKE which only allows range copying (daughter lineages share their ancestral area, Garcia-R & Matzke, 2021). In addition, each of these models was used with a modification to include potential for long-distance (i.e., jump) dispersal (+J) (Matzke, 2014); (4) DIVALIKE+J, (5) DEC+J, and (6) BAYAREALIKE+J (Garcia-R & Matzke, 2021). We used 1,000 randomly sampled post-burn-in trees from the BEAST analyses results as input. We assigned species between Nearctic and Neotropical realms (Fig. 1). The boundary between realms was the Trans-Mexican Volcanic Belt (Rico *et al*., 2022). We used sample-size corrected Akaike weights (AIC_c_) to compare fitness among models (Garcia-R & Matzke, 2021).

**FIGURE 1.**
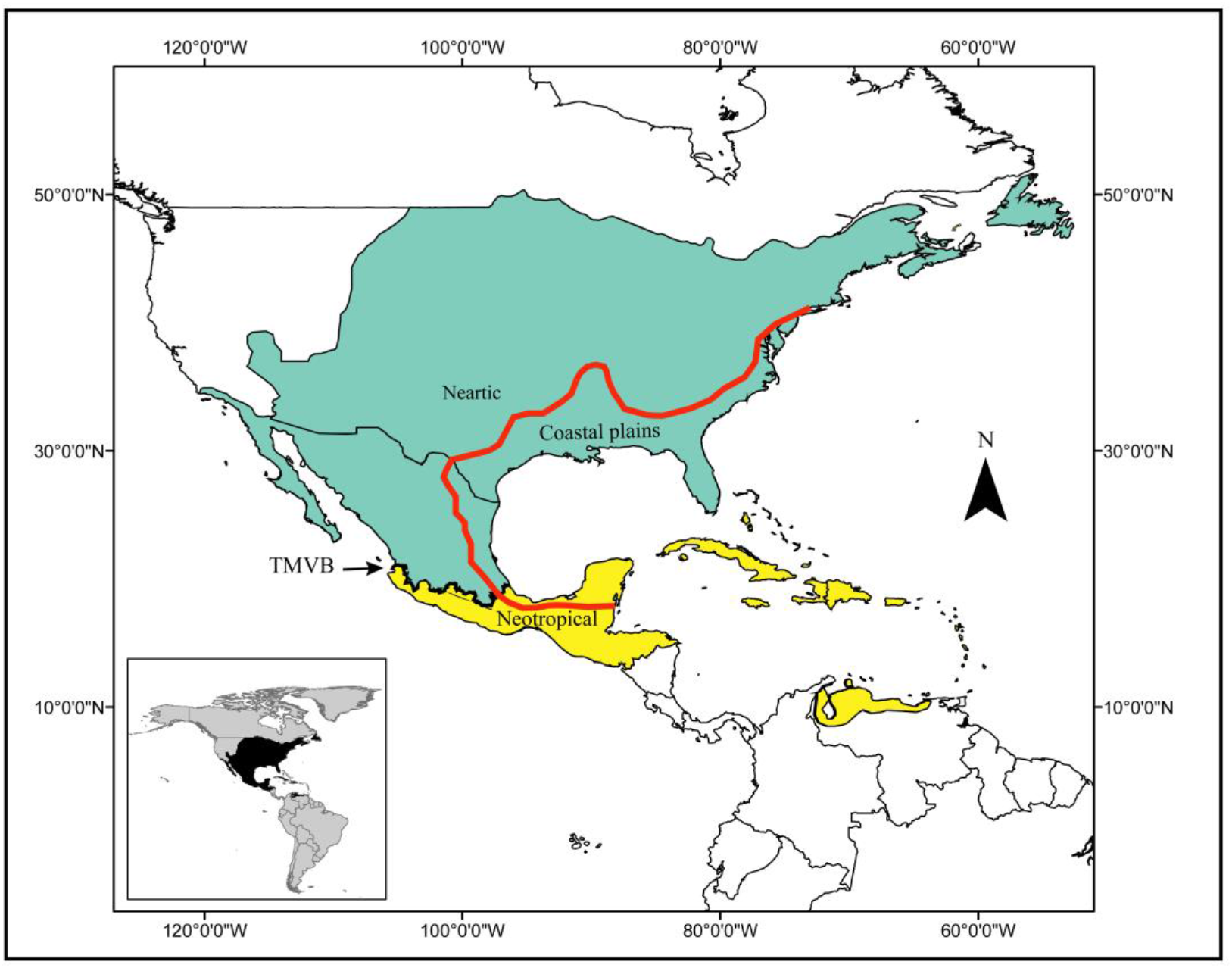
Map depicting the distribution of Funduloidea between North and northern South America. The boundary between realms is the Trans-Mexican Volcanic Belt (TMVB; Rico et al., 2022). The red line depicts the boundary between the coastal plain physiographic province and continental uplands. Inset shows Funduloidea distribution within the Americas.

### Ancestral-habitat reconstruction

To document the evolution of habitat affiliation, we used the make.simmap function in R (R Development Core Team, 2021), using the Phytools package (Revell, 2012). This procedure fits a continuous-time reversible Markov model for the evolution of a character, using a simulated model and tree-tip designations by species to simulate stochastic character histories (Revell, 2012). The procedure was performed with 1000 stochastic character histories simulations, choosing the equal-rates model, which assumes that a trait is acquired or lost under the same probabilities over time (Passarotto *et al*., 2018). Habitat data for each species were obtained from literature, classified as: (1) coastal species occurring inland only within the coastal plain physiographic province (Fig. 1), (2) upland species of inland physiographic regions above the coastal plain, and (3) species with combined coastal-upland distributions (Supporting information, Table S2).

## RESULTS

### Phylogenetic analyses

The Mr. Bayes and BEAST produced similar topologies. Bayesian inference recovered almost all genera with posterior probabilities >0.95 (Fig. 2; Table 2). Most genera were monophyletic except (1) *Xenotoca variata* was sister to *Ameca splendens*, (2) *Girardinichthys turneri* was sister to remaining *Girardinichthys* plus *Neotoca* and *Skiffia*, and (3) *Jordanella pulchra* was sister to *Floridichthys* while *J. floridae* was sister to *Megupsilon-Cyprinodon*. Congruent with genomic results (Ghezelayagh *et al*., 2022; Piller *et al*., 2022), our results indicate Cyprinodontidae is within Funduloidea, sister to Fundulidae (Cyprinodontidae + Fundulidae, PP >0.95) and *Cubanichthys* is sister to Profundulidae + Goodeidae.

**Table 2.** Divergence estimates with confidence intervals (CIs) for numbered chronogram nodes (Fig. 2) that are evaluated in this study. Environmental events potentially associated with phylogenetic divergences (reciprocal illumination) are provided. Further description is provided in text with references.

| Node | Divergence (node) | Age estimate<br>(MA)<br>(95% HPD<br>CI) | Reciprocal illumination |
| --- | --- | --- | --- |
| 1 | Funduloidea split Neotropics / Nearctic | **61.76**<br>**(48.53-76.99)** | 59.2-47.8 Ma – Tropic of Cancer retracted to 27°N (Nearctic-Neotropical subdivision)<br>55.8-55.0 Ma – Gulf of México drawdown (coastal barriers) |
| 2 | Fundulidae / Cyprinodontidae | **55.09**<br>**(45.10-72.34)** | 55.8-55.0 Ma – Gulf of México drawdown (coastal barriers)<br>55.5 Ma – North American drainage reorganization (coastal barriers, niche shift) |
| 3 | Goodeidae-Profundulidae / Cubanichthyidae | **55.94**<br>**(43.07-70.49)** | 66.0-56.0 Ma – Gulf-Caribbean seaway (dispersal between seas)<br>55.8-55.0 Ma – Gulf of México drawdown (continental barriers) |
| 4 | Goodeidae / Profundulidae | **37.76**<br>**(27.81-48.44)** | 40.0-25.0 Ma – Chiapas Massif uplift (inland drainage capture, Profundulidae)<br>38.5-33.9 Ma – Madrean River isolated from Gulf (inland drainage capture, Goodeidae)<br>33.9-27.8 Ma – Tropic of Cancer retracted to 25°N (Nearctic-Neotropical subdivision) |
| 5 | *Profundulus* / *Tlaloc* | **28.33**<br>**(19.65-37.42)** | 35.0-25.0 Ma – Erosion of Chiapas Massif foreland (river capture)<br>30.0-25.0 Ma – Northern Chiapas Massif uplift (river capture) |
| 6 | Empetrichthyinae / Goodeinae | **21.15**<br>**(15.61-27.02)** | 23.0 Ma – North American drainage reorganization, Great Basin tectonism (range fragmentation) |
| 7 | Characodontini-Ilyodontini / Goodeinae | **14.15**<br>**(10.58-17.96)** | 14.0-5.0 Ma – San Marcos Fault reactivation (range fragmentation)<br>13.6-10.6 Ma – Los Encinos volcanism (San Luis Potosí) (range fragmentation)<br>12.0 Ma – Metates Volcanism (Durango) (range fragmentation) |
| 8 | Cyprinodontidae / *Jordanella pulchra*-*Floridichthys* | **39.96**<br>**(28.54-52.27)** | 47.8-33.9 Ma – Middle-Late Eocene cooling (dispersal into tropical Yucatán) |
| 9 | *Cyprinodon*-*Megupsilon*-*Jordanella* / *Cualac* | **26.73**<br>**(18.49-35.66)** | 32.1-17.6 Ma – Sea-level falls (coastal barriers)<br>27.8-16.0 Ma – Late Oligocene-Early Miocene Chicotepec Basin uplift (inland drainage capture) |
| 10 | *Jordanella floridae* / *Megupsilon*-*Cyprinodon* | **19.26**<br>**(12.65-26.20)** | 23.0 Ma – Sea-level fall (dispersal to Ocala High)<br>18.0 Ma – Coastal flooding (island barrier)<br>17.0-13.8 Ma – Middle Miocene Climatic Optimum (island barrier) |
| 11 | *Megupsilon* / *Cyprinodon* | **11.02**<br>**(7.68-14.84)** | 11.8 Ma – Development of Río San Fernando (dispersal inland) |
| 12 | *Cyprinodon* / Yucatán *Cyprinodon* | **7.28**<br>**(5.40-9.30)** | 11.6-5.3 Ma – Uplift of Yucatán Peninsula (continental barrier) |
| 13 | Fundulidae / *Leptolucania* | **43.69**<br>**(33.62-54.22)** | 43.6-40.8 Ma – Sea-level falls (dispersal to Ocala High)<br>40.1 Ma – Sea-level rise (island barrier) |
| 14 | *Fundulus* / *Lucania* | **43.69**<br>**(27.38-42.76)** | 33.9-33.7 Ma – Sea-level fall (coastal barrier) |
| 15 | *Fundulus* / *Wileyichthys* | **31.79**<br>**(25.15-38.92)** | 38.5-23.0 Ma – Ancestral Río Grande (dispersal inland)<br>34.0-28.0 Ma – Volcanism (range fragmentation) |
| 16 | *Fundulus* main split | **26.77**<br>**(21.6-32.1)** | 32.1-17.6 Ma – Sea-level falls (coastal barriers) |
|  |  |  | 26.0 Ma – Hackberry Embayment (coastal barriers) |
| 17 | *Fundulus* / *Fundulus majalis*-*F. persimilis*-*F. similis* | **22.38**<br>**(18.23-23.80)** | 23.0 Ma – Houston-Brazos estuary abandoned (niche shift) |
| 18 | *Zygonectes* / *F. sciadicus*-*Plancterus* | **22.12**<br>**(17.37-27.14)** | 23.0 – Emergence of North American grasslands (niche shift) |

**FIGURE 2.**
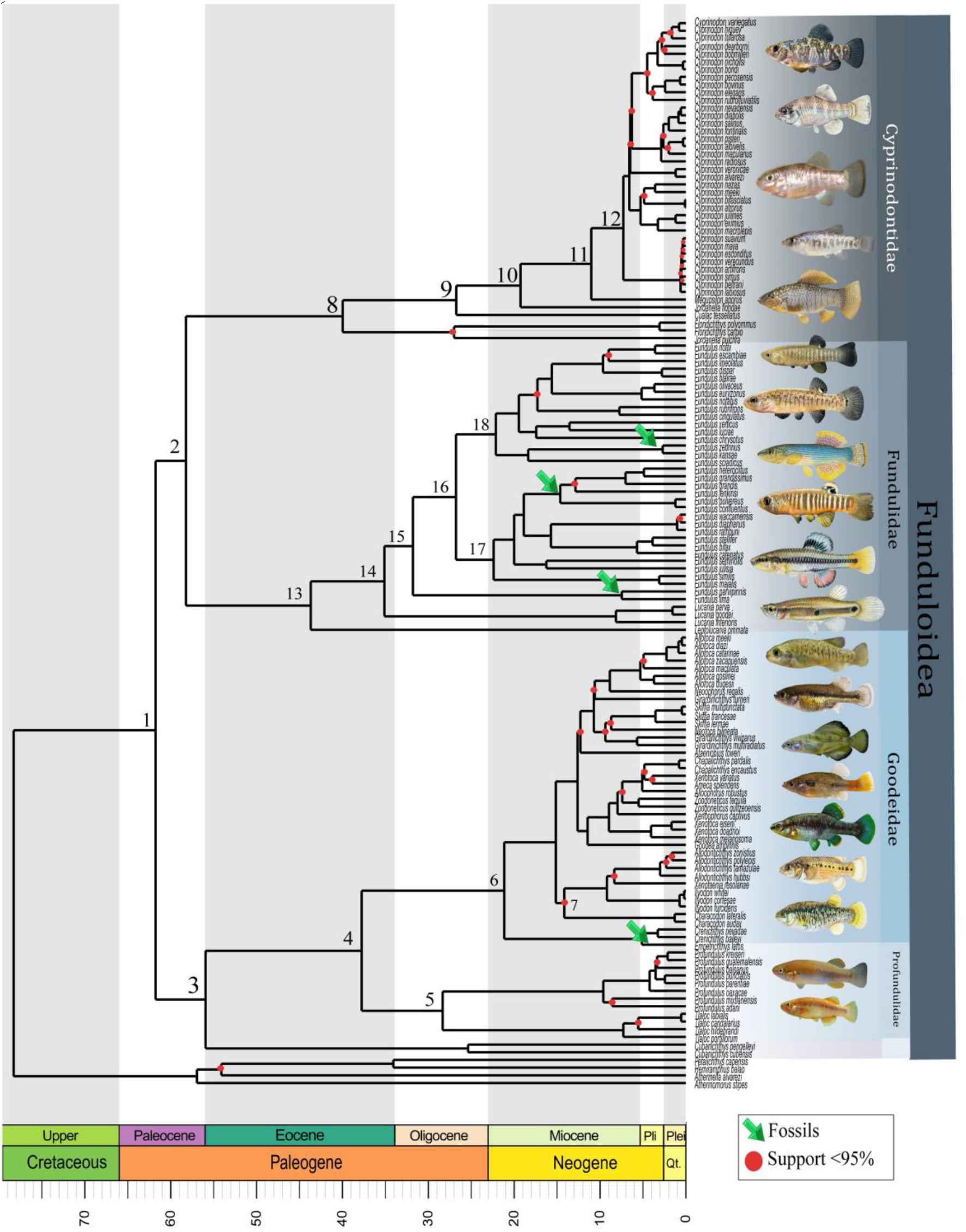
Phylogenetic relationships and divergence times among taxa within Funduloidea. This topology was recovered by Bayesian inference with five concatened loci (three mitochondrial, two nuclear) and 135 species. Red dots depict nodes with posterior probabilities <95%. Green arrows show fossil calibration points (Table 1). Numbered nodes are associated with environmental and geologic events in our biogeographical synthesis (Table 2).

### Diversification-through-time

The LTT plots (Fig. 3), indicate Cyprinodontidae is the only family that experienced a diversification-rate shift, initiating ~11.02 Ma and continuing to present (λ = 0.18 mean; range = 0.08-0.43). Initial rate increase occurred when *Megupsilon* diverged from *Cyprinodon*. No shifts were detected in Fundulidae (λ = 0.07 mean; range = 0.03-0.18), Goodeidae (λ = mean 0.15; range = 0.06-0.38), or Profundulidae (λ = mean 0.12; range = 0.02-0.36).

**Figure 3.**
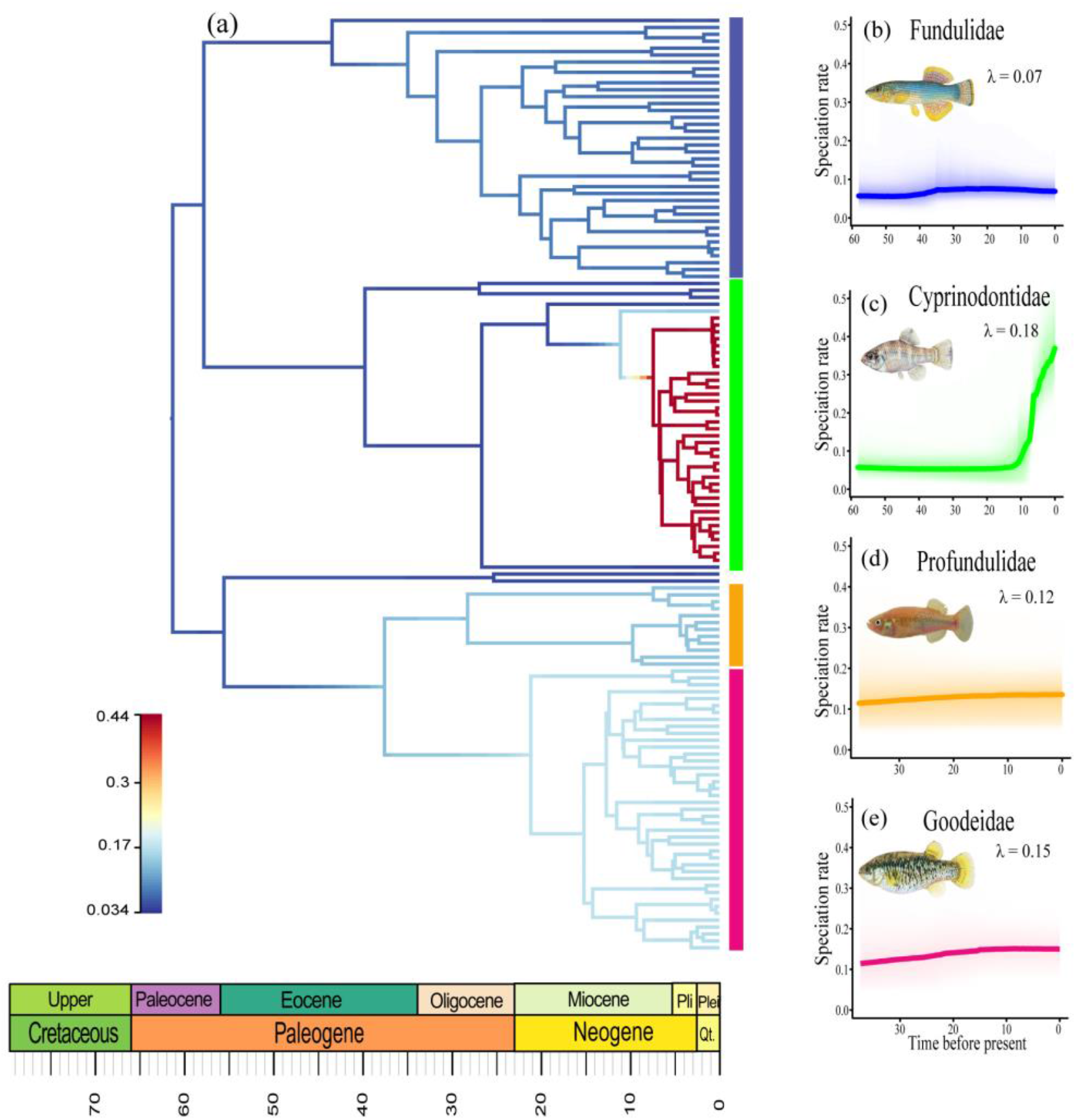
(a) BAMM phylorate plot showing speciation rates along each Funduloidea branch. Cool colors represent slow rates; warm colors represent fast rates. Speciation rates are shown for (b) Fundulidae, (c) Cyprinodontidae, (d) Profundulidae, (e) Goodeidae. λ = mean diversification rate.

### Ancestral-areas reconstruction

The best-fit model for ancestral-areas reconstruction was the BAYAREALIKE+J (Supporting information, Table S3, consistent with the coarse level of analysis, which recognized only the Nearctic and Neotropical realms. Most divergence events occurred within one realm or the other, as required by the BAYAREALIKE model (Garcia-R & Matzke, 2021). Inclusion of the +J portion of the model implies that niche shifts between realms occurred via long-distance dispersal (Matzke, 2014).

The ancestral distribution for Funduloidea is optimized as Nearctic-Neotropical, suggesting a widespread most recent common ancestor (MRCA) (Fig. 4). Funduloidea first split into a northern fork with primarily Nearctic affinities (Fundulidae-Cyprinodontidae) and southern fork with Nearctic-Neotropical affiliation (Cubanichthyidae, Profundulidae, Goodeidae). Appearance of fundulids in the Neotropics is relatively recent (Late Neogene-Quaternary), limited to the *Fundulus grandis* species group. Nearctic Cyprinodontidae made several Neotropical invasions (*Floridichthys*, Yucatán *Cyprinodon*, Caribbean *Cyprinodon, C. variegatus* species group). The MRCAs of Cubanichthyidae and Profundulidae were Neotropical and both families are restricted to that realm. Goodeidae most likely had a Nearctic origin, with representatives of *Girardinichthys, Allodontichthys, Xenotaenia*, and *Ilyodon* reaching Neotropical drainages in the Late Neogene-Quaternary.

**FIGURE 4.**
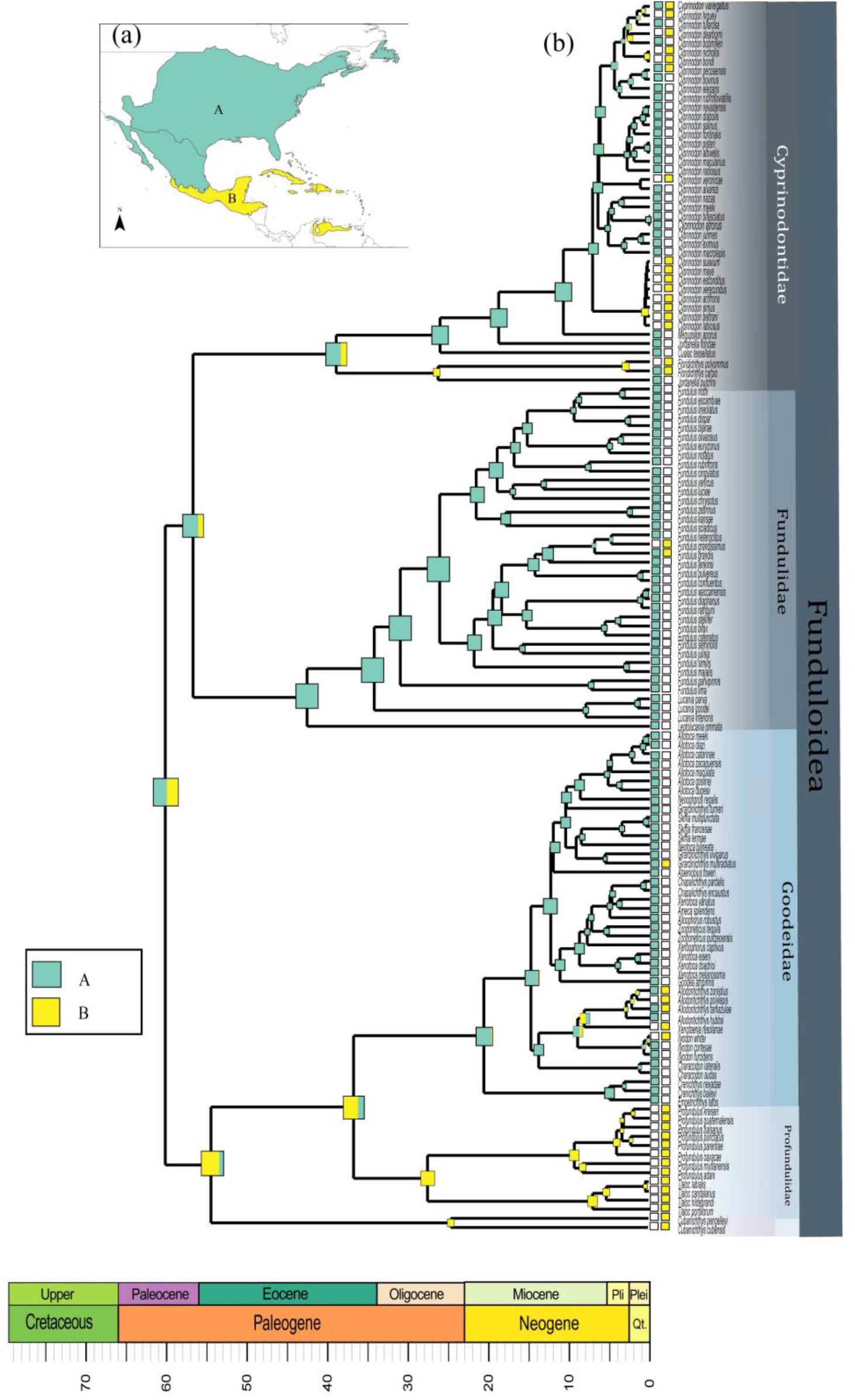
(a) distribution of Funduloidea species divided as Nearctic = A and Neotropical = B, (b) ancestral areas computed using the BAYAREALIKE+J model in BioGeoBEARS.

### Ancestral-habitat reconstruction

The ancestral-habitat reconstruction (Fig. 5) indicates that the MRCA of Funduloidea was coastal. Inland invasions within Fundulidae include *Lucania interioris*, the *F. sciadicus-Plancterus* group, and later-branching lineages within subgenera *Fundulus* and *Zygonectes*. Fundulidae is the only family with species broadly distributed between coastal and upland habitats. In Cyprinodontidae, *Floridichthys* and *Jordanella* retained coastal affinities while the maritime branch of *Cyprinodon* gave rise to upland and coastal lineages. Other branches within *Cyprinodon* became upland lineages. Cubanichthyidae remained coastal, whereas Profundulidae and Goodeidae originated and diversified entirely in uplands.

**FIGURE 5.**
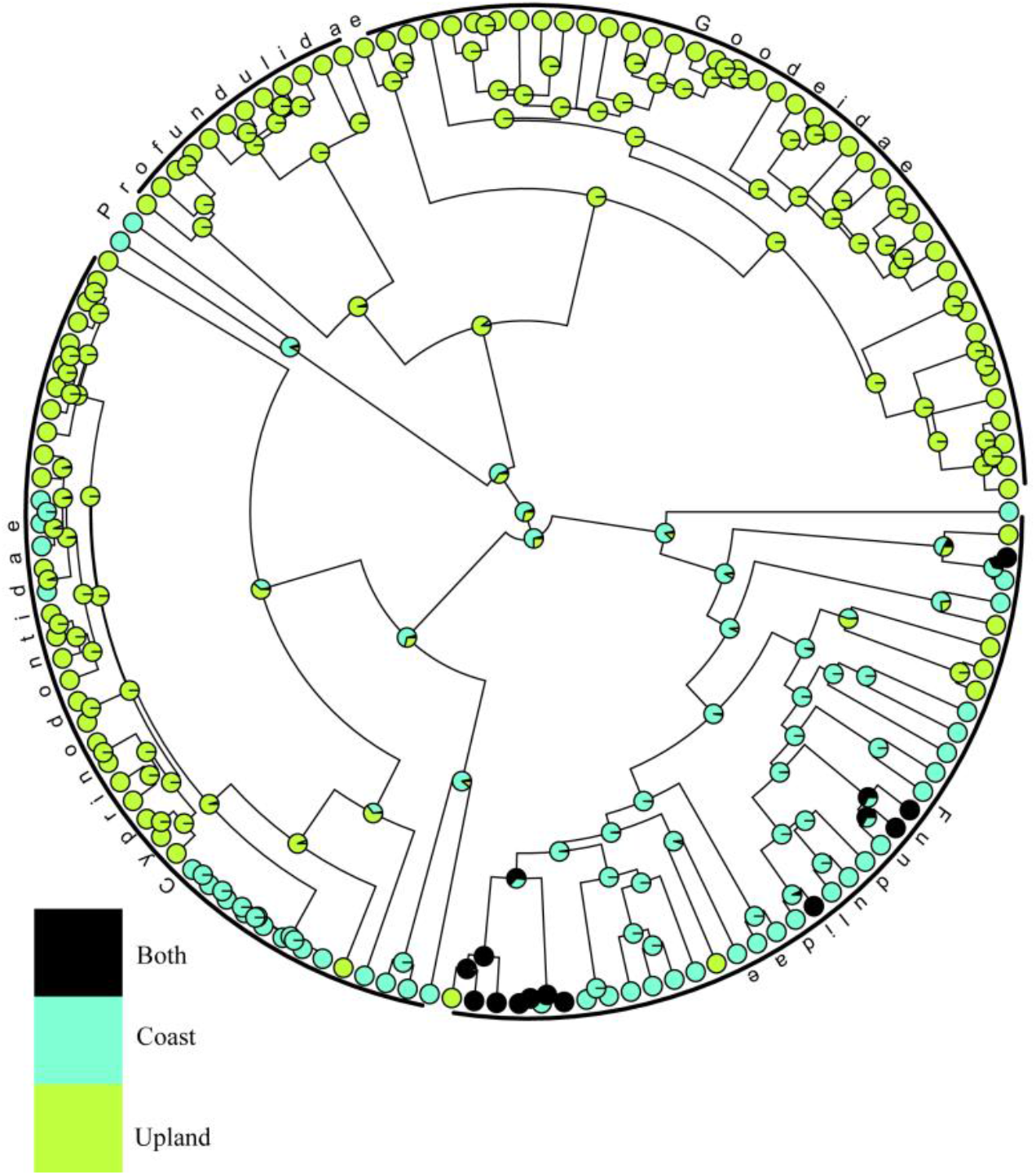
Ancestral-habitat reconstruction. Pie graphs at each node represent the probability of each character state: Coast, Upland, Both.

### Early branching in Funduloidea

Evidence suggests the MRCA of Funduloidea emerged in the Late Cretaceous Gulf of México, spanning present-day Nearctic and Neotropical realms. The MRCA was likely tropical because the tropics extended to 30° North at this time (Zhang *et al*., 2019). The MRCA also likely had high salinity tolerance (Ghedotti & Davis, 2013). Cretaceous origin indicates the MRCA survived the Chicxulub meteor impact ~66 Ma. Given extreme impacts of this event (Schulte *et al*., 2010), it may have been confined to an isolated refugium.

The primary split in Funduloidea, 77.0-48.5 Ma (Fig. 2, node 1), suggests north-south division. The Tropic of Cancer retreated to 27°N by the Early Eocene (Zhang *et al*., 2019), suggesting the Nearctic MRCA of Fundulidae-Cyprinodontidae (Fig. 4) inhabited the northwestern Gulf Coast. A lack of river deltas for several hundred kilometers south of the Río Grande (Snedden & Galloway, 2019) possibly caused a distributional hiatus of funduloids. The MRCA of Cubanichthyidae-Goodeidae-Profundulidae likely populated the next delta to the south, fed by a large drainage extending northwest along the front of the Sierra Madre (Snedden & Galloway, 2019; henceforth, Madrean River). Evidence suggests the Gulf of México became isolated from the world ocean 55.80-54.95 Ma, accompanied by drawdown 900-1300 m (Cossey *et al*., 2016, 2021). Because sea-level fall isolates estuaries (Dolby *et al*., 2020), this event could have separated northern Fundulidae-Cyprinodontidae from southern Cubanichthyidae-Goodeidae-Profundulidae (Table 2).

Divergence of Cyprinodontidae from Fundulidae, 72.3-45.1 Ma (Fig. 2, node 2), may also result from Gulf of México drawdown (Table 2) along with a ~55.5 Ma reconfiguration of drainages feeding the Gulf of México (Snedden & Galloway, 2019). We suggest ancestral Fundulidae populated the Houston-Brazos River delta, the only fluvial delta then present (Snedden & Galloway, 2019). In contrast, ancestral Cyprinodontidae putatively originated from an ancestor like *C. variegatus* (Echelle & Echelle, 2020), a species well adapted for open coasts with fluctuating temperatures and salinities (Nordlie, 2006). We infer this taxon inhabited the Rosita Delta, which received diffuse freshwater inflow, but was wave-dominated and sandy (Snedden & Galloway, 2019). Gulf drawdown potentially produced extreme salinities (Cossey *et al*., 2016, 2021), perhaps explaining exceptional tolerance of fluctuating salinities among cyprinodontids.

Divergence of Cubanichthyidae from Goodeidae-Profundulidae, 70.5-43.1 Ma (Fig. 2, node 3), could also link to the Gulf drawdown (Table 2). A Paleocene seaway along the Mexican foreland and across the Yucatán platform (Martens & Sierra-Rojas, 2021) potentially allowed ancestral *Cubanichthys* to reach the Caribbean Sea. Gulf drawdown would have dewatered the seaway, isolating *Cubanichthys* to the east.

Divergence of Goodeidae from Profundulidae, 48.4-27.8 Ma (Fig. 2, node 4), presumably reflects upland isolation of lineages in separate river drainages (Table 2, Fig. 4). The ancestral goodeid likely occupied the Madrean River (Río Bravo in Snedden & Galloway, 2019), which was cut off from the Gulf of México in the Late Eocene (Galloway *et al*., 2011). By the end of the Eocene, the tropics retreated to 25°N (Zhang *et al*., 2019), making the Madrean River Nearctic. For Profundulidae, the timing of divergence corresponds with uplift of the Chiapas Massif (40-25 Ma, Villagómez & Pindell, 2020). Drainage from the Massif to the Bay of Campeche was localized at this time, not reaching the Gulf of México basin (Beltrán-Triviño *et al*., 2021; Villagómez *et al*., 2022).

### Profundulidae

The major node within Profundulidae, 37.4-19.7 Ma (Fig. 2, node 5), separates *Profundulus* from *Tlaloc. Tlaloc* occurs across the Chiapas Massif and adjacent highlands within the Río Grijalva drainage (Miller, 1955; Cashner & Echelle, 2020). Upper Río Grijalva initially flows northwest but makes an abrupt turn northeast, a pattern consistent with capture by a north-flowing river, possibly the ancestral Río Uxpanapa or Tonalá, caused by uplift of the northwestern Chiapas Massif 30-25 Ma (Witt *et al*., 2012), or head-cutting of the lower Río Grijalva during a period of extensive erosion 35-25 Ma (Abdullin *et al*., 2016). Divergence of *Tlaloc* from *Profundulus* provides an estimate for timing of this capture (Table 2) and explains the distinctiveness of the Upper Grijalva fish community (Elías *et al*., 2020). Ancestral *Profundulus* presumably remained west of the Chiapas Massif (ancestral Río Coatzacoalcos). Rising sea levels could explain disappearance of *Profundulus* from Atlantic-slope drainages, as proposed for *Herichthys* (Pérez-Miranda et al., 2020). During the Middle Miocene Climatic Optimum (MMCO), 17.0-13.8 Ma (Miller *et al*., 2020a), seas inundated areas north and east of the Sierra Madre del Sur and Chiapas Massif (Blakey & Ranney, 2018). Diversification occurred thereafter, possibly from one refugial population.

### Goodeidae

Oligocene transfer of the upper Madrean River to the Río Grande (Galloway *et al*., 2011; Snedden & Galloway, 2019) allowed Goodeidae to expanded westward, along the front of the Sierra Madre Occidental, to the Great Basin. Divergence of Empetrichthyinae 27.0-15.6 Ma (Fig. 2, node 6, Table 2) agrees with breakup of this drainage at the Oligocene-Early Miocene transition (Snedden & Galloway, 2019). Great distance between northern (Empetrichthyinae) and southern (Goodeinae) goodeids is a biogeographic anomaly (Webb, 2020). Parenti (1981) proposed regional desiccation eliminated intervening populations (see also Grant & Riddle, 1995; Webb *et al*., 2004; Webb, 2020), but living Empetrichthyinae occupy the driest region of North America (i.e., aridity is associated with survival). Fragmentation of the Madrean River may have been an additional cause of extirpations (Fagan *et al*., 2002) and climatic cooling may have reduced habitat suitability at higher elevations and latitudes, perhaps explaining why Empetrichthyinae associates with warmwater springs. It is also possible remnant goodeid populations were present in the uninhabited region during European settlement, but disappeared due to habitat destruction, dewatering, or invasive species associated with European contact. Because this scenario agrees with reconstructed hydrography, it is unnecessary to invoke the popular hypothesis (Minckley *et al*., 1986; Grant & Riddle, 1995; Miller *et al*., 2005; Webb, 2020) that northward drift of the Pacific Plate created the gap between Empetrichthyinae and Goodeidae.

The hypothesis that Goodeidae used a western route from the Great Basin into central México (Pérez-Rodríguez *et al*., 2015) requires modification because Goodeidae dispersing from the Gulf of México would have reached central México en route to the Great Basin. Complex geological history confounds understanding of Middle Miocene hydrology, but reactivation of the San Marcos Fault, 14-5 Ma and Middle Miocene volcanism on the northwest and eastern borders of the Mesa Central (Aranda-Gómez *et al*., 2007; Chávez-Cabello *et al*., 2007; Nieto-Samaniego *et al*., 2007), potentially disrupted the Madrean River. Divergence of Characodontini-Illyodontini, 18.0-10.6 Ma (Fig. 2, node 7), appears to have been an east-west divergence (possibly between ancestral ríos Nazas and Aguanaval, Table 2). Loss of Goodeids from the Gulf of México drainage could reflect Late Miocene integration of the ancestral Río Nazas with the Río Grande, which facilitated an influx of fishes (Hoagstrom & Osborne, 2021). *Characodon* persisted only in remote basins (†*C. garmani*, Mayran-Parras Basin; *C. audax-C. lateralis*, Río Tunal (Beltrán-López *et al*., 2021), potentially protected from invading fishes. Positioning of Goodeinae on the TMVB throughout its tectonic evolution, which lasted 17 my (Ferrari *et al*., 2012), the lineage experienced frequent drainage reorganizations and shifting hydrographic barriers (Barbour, 1973; Domínguez-Domínguez *et al*., 2010; Pérez-Rodríguez *et al*., 2015). Stream capture was likely the main mode of range expansion and diversification (Webb *et al*., 2004; Domínguez-Domínguez *et al*., 2010; Beltrán-López *et al*., 2021).

### Cyprinodontidae

From Middle to Late Eocene, the estuary of the Madrean River disappeared, a wave-dominated shore zone developed (Snedden & Galloway, 2019), and the climate cooled (Zachos *et al*., 2001; Miller *et al*., 2020b). These conditions potentially favored southward invasion by Cyprinodontidae, following the into-the-tropics paradigm (Meseguer & Condamine, 2020). Divergence of *Jordanella pulchra*-*Floridichthys*, 52.3-28.5 Ma (Fig. 2, node 8, Table 2), implies Cyprinodontidae inhabited the southwestern Gulf by the Late Eocene. Possibly, this invasion displaced coastal Goodeidae and Profundulidae, leaving only inland populations discussed above.

Because living *Floridichthys* have a disjunct distribution across the Gulf of México, the MRCA could be from Florida or Yucatán. Presence of the sister taxon *Jordanella pulchra* in Yucatán favors that scenario and is compatible with the hypothesis (above) that Cyprinodontidae originated in the Early Eocene Rosita Delta (closer to Yucatán than Florida). Further, the Caribbean-Loop Current could have facilitated later oceanic dispersal of *Floridichthys* from Yucatán to Florida. Hence, we suggest the Late Eocene ancestor of *J. pulchra-Floridichthys* dispersed along the western Gulf Coast to the Bay of Campeche where coastal and shallow-marine habitats (Villagómez *et al*., 2022) provided suitable habitats. However, *J. pulchra-Floridichthys* has uncertain phylogenetic placement (compare Fig. 2 with Piller *et al*., 2022), so further study is needed.

Endemism of *Cualac tesselatus* in the Río Pánuco, 35.7-18.5 Ma (Fig. 2, node 9), implies the MRCA inhabited the western Gulf Coast in the Oligocene and ancestral *C. tesselatus* was living within the ancestral Río Pánuco (mapped in Beltrán-Triviño *et al*., 2021). Late Oligocene-Early Miocene uplift of the Chicontepec Basin (Roure *et al*., 2009), combined with Oligo-Miocene sea-level falls roughly every 1.2 my (Oi2-Mi1 low stands, 32.1-17.6 Ma, Boulila *et al*., 2011), may have stranded this population inland (Table 2).

Our phylogeny indicates *Jordanella* is polyphyletic. Separation of *Jordanella floridae* from *Megupsilon*-*Cyprinodon*, 26.2-12.7 Ma (Fig. 2, node 10), suggests an ancestral cyprinodontid dispersed eastward during the Early Miocene. The Florida Peninsula was and island (i.e., Ocala High) and lowered sea levels reduced the marine gap with the mainland (Popenoe, 1990), possibly facilitating continent-to-island immigration (Table 2). Subsequent coastal flooding 18 Ma (Snedden & Galloway, 2019) and elevated MMCO sea-levels (17.0-13.8 Ma, Miller *et al*., 2020) could have isolated ancestral *J. floridae* (Table 2).

Late Miocene divergence of *Megupsilon* (14.8-7.7 Ma, Fig. 2, node 11) may represent Late Miocene emergence of ancestral Río San Fernando (Río Bravo in Snedden & Galloway, 2019) as a corridor for inland invasion (Table 2). This species (extinct in the wild) inhabited a springfed system (Miller *et al*., 2005), and may have speciated as a spring endemic. It is possible episodes of aridity, tectonism, or volcanism isolated the spring system, but this needs further study.

Yucatán *Cyprinodon* descend from the first diverging lineage of the genus (9.3-5.4 Ma, Fig. 2, node 12). Emergence of the peninsula in the Late Miocene (Bautista *et al*., 2011) was contemporary with this divergence (Table 2) and with climatic cooling (Zachos *et al*., 2001). This scenario agrees with the into-the-tropic hypothesis (Meseguer & Condamine, 2020).

Rate of *Cyprinodon* speciation increased upon separation from *Megupsilon* (node 11, Fig. 2-3). Several factors likely contributed to this trend. First, inland invasions into five Late Miocene rivers subdivided *Cyprinodon* into as many upland lineages, four of which dispersed across the desert region (Echelle *et al*., 2005; Hoagstrom & Osborne, 2021). Once inland invasions were underway, drainage rearrangements, climate fluctuations, tectonism, and volcanism caused widespread allopatric diversification (Echelle, 2008; Knott *et al*., 2008; Hoagstrom & Osborne, 2021). Meanwhile, a maritime lineage of *Cyprinodon* remained along the Gulf Coast (Echelle *et al*., 2005, 2006), which is unclear in our ancestral-habitat reconstruction (Fig. 5) because the same widespread ancestor produced each upland invasion while remaining on the coast. During the Pleistocene, maritime *Cyprinodon* made new invasions into the desert region (Lozano-Vilano & Contreras-Balderas, 1999; Hoagstrom & Osborne, 2021) and immigrated to Caribbean islands and South America (Haney *et al*., 2007, 2009). The Yucatán lineage produced a species flock (Strecker, 2006) (Fig. 3). Beyond this, our analysis may underestimate the *Cyprinodon* speciation rate because a species flock from San Salvador, Bahamas (Martin & Wainwright, 2013) and *C. variegatus hubbsi* (a potential distinct species, Jung *et al*., 2019) are absent from our phylogeny, which also leaves out seven *Cyprinodon* species for which we had no genetic data. Finally, six of the recognized species included in the analysis are likely polyphyletic (Echelle & Echelle, 2020), but represented as one taxon here.

### Fundulidae

Early Eocene fundulids were theoretically living in the delta of the Houston-Brazos River. Divergence of *Leptolucania*, 54.2-33.6 Ma (Fig. 2, node 13), suggests immigration across the Suwannee Channel, to the Ocala High (Table 2), which developed peritidal landforms at this time (Avon Park and Ocala formations, Randazzo & Jones, 1997; Maliva *et al*., 2011). Sea-level falls of 10-20 m at 43.6, 42.9, and 40.8 Ma (Miller *et al*., 2020a) could have facilitated immigration. Subsequent sea-level rise (40 m, Middle Eocene Climatic Optimum, 40.1 Ma) could have isolated Ocala populations. Although *Leptolucania* is a freshwater genus, the ancestor likely had high salinity tolerance (Ghedotti & Davis, 2013).

Contrary to earlier reconstructions (Ghedotti & Davis, 2017), our phylogeny indicates *Lucania* is sister to *Fundulus*. As *Lucania interioris* is the earliest diverging lineage, we hypothesize *Lucania* originated in the northwestern Gulf of México. If high sea levels of the Middle Eocene allowed the MRCA of *Fundulus-Lucania* to expand southwest along the Gulf Coast, sea-level fall ~75 m (33.9-33.7 Ma, Miller *et al*., 2020) could have isolated southwestern populations in the conjoined Río Grande-Río San Fernando delta (Snedden & Galloway, 2019), in agreement with divergence of *Lucania*, 42.8-27.4 Ma (Fig. 2, node 14, Table 2).

Divergence of *Fundulus* subgenus *Wileyichthys* 38.9-25.2 Ma (Fig. 2, node 15, Table 2), the MRCA invaded the ancestral Río Grande, which extended northwest to the Continental Divide, proximate to the Pacific Coast (Blakey & Ranney, 2018; Snedden & Galloway, 2019). Fossil *Fundulus* in the Great Basin (Smith *et al*., 2002) are consistent with this hypothesis. A pulse of intense volcanism in the northern Sierra Madre Occidental 34-28 Ma (Ferrari *et al*., 2018) presumably isolated *Wileyichthys* from relatives in the Gulf of México watershed.

The major split within *Fundulus*, 32.1-21.6 Ma (Fig. 2, node 16) corresponds to sea-level oscillations every 1.2 Ma (Oi2-Mi1 low stands, 32.1-17.6 Ma, Boulila *et al*., 2011), which reached 30-40 m below modern sea level (Miller *et al*., 2020b), potentially separating the Mississippi and Houston-Brazos deltas (Table 2). Also, ~26 Ma, a slope failure west of the Mississippi Delta created the Hackberry Embayment (Snedden & Galloway, 2019), which might have been a deepwater barrier between deltas. We hypothesize *Zygonectes-F. sciadicus-Plancterus* originated in the Mississippi River delta because *Plancterus* (node 18) is Mississippian and is the westernmost lineage on this branch. By default, subgenus *Fundulus* presumably originated in Houston-Brazos River delta. Later abandonment of the Houston-Brazos Delta (Oligo-Miocene transition, 23 Ma, Snedden & Galloway, 2019) could have had transferred subgenus *Fundulus* into the new Red River delta, which later merged with the Mississippi (15 Ma, Snedden & Galloway, 2019), potentially bringing subgenus *Fundulus* along.

### Subgenus *F**undulus*

Earliest branching within subgenus *Fundulus* reflects ecological speciation. *Fundulus majalis-F. persimilis-F. similis* inhabit unvegetated coastal habitats where they dive into soft sediments for cover rather than retreating to vegetation like typical *Fundulus* (Martin & Finucane, 1968; Harvey, 1998). They are adapted for continuous swimming in the surf zone (Yetsko & Sancho, 2015) and spawn in unvegetated habitats (Greeley *et al*., 1986). *Fundulus majalis* segregates from congeners in unvegetated, high salinity waters (Weisberg, 1986; Wagner & Austin, 1999). We propose this lineage represents an ancestor adapted for open coasts. Divergence 23.8-18.2 Ma (Fig. 2, node 17) was concurrent with abandonment of the Houston-Brazos delta, suggesting this lineage might represent populations left behind and adapted to surf-zone habitats that developed once the Houston-Brazos delta was abandoned.

### Subgenus *Z**ygonectes**-F*. *sciadicus**-P**lancterus*

Our Funduloidea tree uniquely groups *F. sciadicus* as sister to *Plancterus* (*F. kansae-F. zebrinus*). Divergence from *Zygonectes*, 27.1-17.4 Ma (Fig. 2, node 18, Table 2), suggests the MRCA of *F. sciadicus-Plancterus* immigrated up the Mississippi River (Fig. 5), which extended to the Rocky Mountains (Snedden & Galloway, 2019). *Fundulus sciadicus* and *Plancterus* are grassland associates (Fausch & Bestgen, 1997), suggesting their MRCA adapted to grassland habitats, which developed across interior North America at this time (Andermann *et al*., 2022). The fossil †*F. detillae* (5.0-2.6 Ma, Ghedotti & Davis, 2017), confirms the lineage occupied the plains.

## DISCUSION

We used evidence from our analyses with literature review to develop a biogeographical scenario of early diversification within Funduloidea. We applied the expectation of divergence through allopatric reproductive isolation, which is predominant for freshwater fishes (Seehausen & Wagner, 2014). We inferred cases of ecological reproductive isolation only when there was no straightforward evidence for allopatry and, at the same time, there was evidence for ecological differentiation. For coastal ancestors, we expected sea-level rise to facilitate range expansion along coastlines, with sea-level falls causing population fragmentation (Dolby *et al*., 2018, 2020). For ocean gaps, we expected the reverse, lowered sea levels should facilitate cross-gap dispersal. We expected periods of climatic cooling to facilitate temperate species invading tropical habitats (into-the-tropics hypothesis, Meseguer & Condamine, 2020). Finally, our scenario maintained geographical consistency and compatibility across the phylogeny (Hoagstrom & Echelle, 2022).

Adaptive radiation (Martin & Wainwright, 2011), viviparity (Helmstetter *et al*., 2016), and transition to freshwater (Bloom *et al*., 2013) may increase speciation rate. Adaptive radiation is observed in *Cyprinodon* (Martin & Wainwright, 2011; Hernandez *et al*., 2018), helping explain accelerated speciation. *Cyprinodon* radiations appear to be associated with interbreeding among lineages establishing secondary contact (Richards *et al*., 2021), suggesting other families lacked genetic potential.

Viviparity is absent in Cubanichthyidae, Cyprinodontidae, Fundulidae, and Profundulidae and we did not detect an increase in speciation rate with viviparity in Goodeidae, in contrast to Helmstetter *et al*. (2016), who placed the origin of Goodeinae at ~8.9 Ma (vs. ~21.2 Ma here). Differences in our methods were (1) we evaluated a species-level phylogeny (rather than genus-level), (2) we partitioned data by gene instead of codon position, (3) we used combined tip- and node-based fossil calibration, and (4) we did not use “†*Cyprinodon breviradius*” for calibration. Other studies (Webb *et al*., 2004; Pérez-Rodríguez *et al*., 2015; Rabosky *et al*., 2018) recovered divergence estimates for Goodeinae relatively similar to ours. Further, the biogeography of Goodeinae is attributable to range expansion and fragmentation during development of the TMVB (Webb *et al*., 2004; Domínguez-Domínguez *et al*., 2010; Pérez-Rodríguez *et al*., 2015), which initiated ~20 Ma (Ferrari *et al*., 2012). A goodein lineage originating only 8.9 Ma (as in Helmstetter *et al*., 2016) would have missed this history and lacked dispersal routes to facilitate its present range.

Transition to freshwater occurs in all major funduloid families but only relates to an increased rate of speciation in Cyprinodontidae. This is interesting for Goodeidae, which also developed viviparity (discussed above), but did not achieve accelerated speciation. Using LTT plots, Pérez-Rodríguez *et al*. (2015) found an early acceleration in speciation, whereas we did not and the slope of our LTT plot resembles a horizontal line (lambda = 0.15 mean). In contrast, transition to freshwater is associated with accelerated speciation in *Cyprinodon* (described above). Poeciliid fishes also exhibit increased diversification within the North American desert (García-Andrade *et al*., 2021) where they cooccur with *Cyprinodon* (Minckley *et al*., 1991), suggesting their shared histories are responsible for their accelerated rates of speciation.

Barrier displacement refers to processes like mountain uplift and river capture, which separate and merge faunas (Albert *et al*., 2017). This process leaves a phylogenetic signal, present in Funduloidea, in which early-branching lineages are geographically peripheral, with limited diversification (Table 3). Most striking within Funduloidea are four sequential instances in Cyprinodontidae, with early-branching lineages isolated south or east of the main distribution of the family. Detailed examples illustrate barrier displacement within Funduloidea (Echelle, 2008; Domínguez-Domínguez *et al*., 2010; Hoagstrom & Osborne, 2021). As a subtle form of barrier displacement, evidence implies there were periodic coastal immigrations by Fundulidae and Cyprinodontidae from the northern Gulf Coast to Yucatán, suggesting populations took advantage of climatic cooling. Relyea, (1983) inferred that the northern limit of mangroves was an ecological boundary for Nearctic funduloids. If so, periods of cooling may have diminished ecological resistance, giving Nearctic lineages opportunities to invade southward (into-the-tropics hypothesis, Meseguer & Condamine, 2020).

**Table 3.** Phylogenetic evidence for diversification by barrier displacement, early diverging lineages with peripheral geography and relatively limited diversification. Sister clades are provided for comparison. The chronogram node (Fig. 2) is given for each sister relation. ^1^count includes described species missing from our phylogeny (Fig. 2). ^2^Count includes species flock in Chichancanab.

| No | Geographically peripheral, early diverging lineage | Species<sub>1</sub> | Geography | Sister lineage | Species<sub>1</sub> | Geography |
| --- | --- | --- | --- | --- | --- | --- |
| 6 | Empetrichthyinae | 4 | Mojave Desert | Goodeinae | 46 | Trans-Mexican Volcanic Belt |
| 7 | Characodontini-Ilyodontini | 12 | Sierra Madre Occidental | Goodeinae (remainder) | 34 | Trans-Mexican Volcanic Belt |
| 8 | *Jordanella pulchra*-*Floridichthys* | 3 | Yucatán Peninsula | Cyprinodontidae (remainder) | 42 | Northwestern Gulf of México |
| 9 | *Cualac* | 1 | Río Pánuco | Cyprinodontidae (remainder) | 41 | Northwestern Gulf of México |
| 10 | *Jordanella floridae* | 1 | Florida Peninsula | Cyprinodontidae (remainder) | 40 | Northwestern Gulf of México |
| 11 | *Megupsilon* | 1 | Río San Fernando | *Cyprinodon* | 39 | Northwestern Gulf of México |
| 12 | Yucatán *Cyprinodon*<sup>2</sup> | 8 | Yucatán Peninsula | *Cyprinodon* (remainder) | 31 | Northwestern Gulf of México |
| 13 | *Leptolucania* | 1 | Florida Peninsula | *Lucania-Fundulus* | 40 | North-central Gulf of México |
| 15 | *Wileyichthys* | 2 | Pacific Coast | *Fundulus-Plancterus-Zygonectes* | 35 | North-central Gulf of México |
| 18 | *Fundulus sciadicus*-*Plancterus* | 3 | Northwestern plains & prairies | *Zygonectes* | 13 | Southeastern woodlands, prairies, swamps |

Fundulidae is the only funduloid family that diversified mostly within a tectonically stable regional, where rivers generate species diversity through high rates of river captures and low rates of extinction (Albert *et al*., 2018). Drainage rearrangements on the Gulf Coastal Plain likely facilitated diversification within *Fundulus*. In addition, north-south tributaries to the Gulf of México provided glacial refugia (Whitehead, 2010).

## CONCLUSIONS

Although scenarios inferred here deserve further study, the phylogeny of Funduloidea supports environmental evidence, providing reciprocal illumination. For example, sea-level falls coincide in time with likely immigrations to the Florida Platform and with periods of lineage isolation along coastlines. Ancient river courses potentially link inland lineages that are now disjunct. Niche shifts coincide with availability of novel ecosystems (open coasts, grasslands). Evidence from Funduloidea is compatible the Gulf Drawdown hypothesis, which remains under investigation (Cossey *et al*., 2021).

All things considered, Funduloidea diversification can be credited to prolonged persistence in a dynamic region with sufficient time to diversify, ability to diversify along coastlines via climatic and sea-level fluctuations, and ability to invade inland and become broadly dispersed and diversified via barrier displacement. An overall signal of diversification via time-for-speciation agrees with recent studies (Rabosky, 2020; Miller, 2021). We do not detect a freshwater-fish paradox. Diversification occurred at similar rates between coastal and freshwater lineages and accelerated speciation in Cyprinodontidae included freshwater and marine lineages. Funduloidea displays exceptional ability to transition from coastal to inland habitats and appears to illustrate that a tolerant, explorative lineage, surviving over more than sixty-million years, can exploit rare opportunities for range expansions and niche shifts, leading to lineage branching and sustained or even accelerated diversification. Because the Gulf of México has offered many such opportunities throughout history, funduloid killifishes have dramatically diversified, writing their own version of the Gulf’s history in their phylogeny.

## SUPPORTING INFORMATION

Additional Supporting Information may be found in the online version of this article at the publisher’s web-site.

**Table S1** GenBank accession numbers for 139 species used in phylogenetic analyses. One hundred and thirty five species in the ingroup and four species in the outgroup.

**Table S2** Habitat clasification by species. Species assigned to habitat (Upland, Coast and Both) for ancestral habitat reconstruction analysis.

**Table S3** Ancestral area reconstruction models. Models tested in BioGeoBEARS, implemented in RASP. The best-fit model showed in bold and highlighted in green.

**File S1** Molecular data sequence aligment, including two nuclear genes (RAG1, Glyt) and three mitochondrialgenes (Cytb, COI and ND2).

